# Regulatory mechanisms of the dynein-2 motility by post-translational modification revealed by MD simulation

**DOI:** 10.1101/2022.02.17.480877

**Authors:** Shintaroh Kubo, Khanh Huy Bui

**Author notes:** Corresponding authors: Shintaroh Kubo, Department of Anatomy and Cell Biology, McGill University, Montréal, Québec H3A 0C7, Canada., Khanh Huy Bui, Department of Anatomy and Cell Biology, McGill University, Montréal, Québec H3A 0C7, Canada.

## Abstract

Intraflagellar transport for ciliary assembly and maintenance is driven by dynein and kinesins specific for the cilia. It has been shown that anterograde and retrograde transports run on different regions of the doublet microtubule, i.e., separate train tracks. However, little is known about the regulatory mechanism of this selective process. Since the doublet microtubule is known to display specific post-translational modifications of tubulins, i.e. “tubulin code”, for molecular motor regulations, we investigated the motility of ciliary specific dynein-2 under different post-translational modification by coarse-grained molecular dynamics. Our setup allows us to simulate the stochastic stepping behaviors of dynein-2 on un-modified, detyrosinated, poly-glutamylated and poly-glycylated microtubules in silico. Our study revealed that poly-glutamylation can play an inhibitory effect on dynein-2 motility. Our result indicates that poly-glutamylation of the B-tubule of the doublet microtubule can be used as an efficient means to target retrograde intraflagellar transport onto the A-tubule.

## Introduction

Intracellular cargo transport driven by molecular motors along microtubule (MT) tracks is an essential cellular process for the maintenance of homeostasis within the cell. Kinesins and dyneins carry cargos or even organelles in the opposite directions, towards the plus end and minus end of the MT respectively. Intracellular transport can be regulated in many different ways. Different MT associate proteins (MAPs) can act as effectors binding to MT to block or promote a certain types of motor proteins (1). For example, kinesins are inhibited by tau binding on microtubules while dyneins are inhibited by tau at a lesser extent (2).

Alternatively, the cell can regulate transport through the “tubulin code”, in which, tubulins, the core component of MT, undergo a variety of post-translational modifications (PTMs) depending on the cellular or tissue locations (3). PTMs, in turn, can modulate the motility of molecular motors directly or indirectly through the binding of MAPs. While there are many kinds of PTMs found on tubulins, the most common and studied PTM are acetylation, poly-glutamylation, poly-glycylation and tyrosination/detyrosination (3). Most of these PTMs are localized on the C-terminal tails (CTT) of α- and β-tubulins, the regions that are flexible and highly negatively charged (Fig. 1a). The CTT frequently interacts with MAPs. In vitro, kinesin-2 and centromere-associated protein E (CENPE), a kinesin-7 have increased motility and greater processivity on detyrosinated chimeric yeast tubulins (4). On the other hand, tyrosination or detyrosination of MT does not affect the binding and motility of yeast dynein-1. Human kinesin-2 express sensitivity to different length of glutamate residues added to tubulin tail by chemical addition while dynein does not show any difference (4). In mouse sperm flagella, tubulin glycylation deficiency leads to abnormal axonemal dynein activity (5).

**Figure 1.**
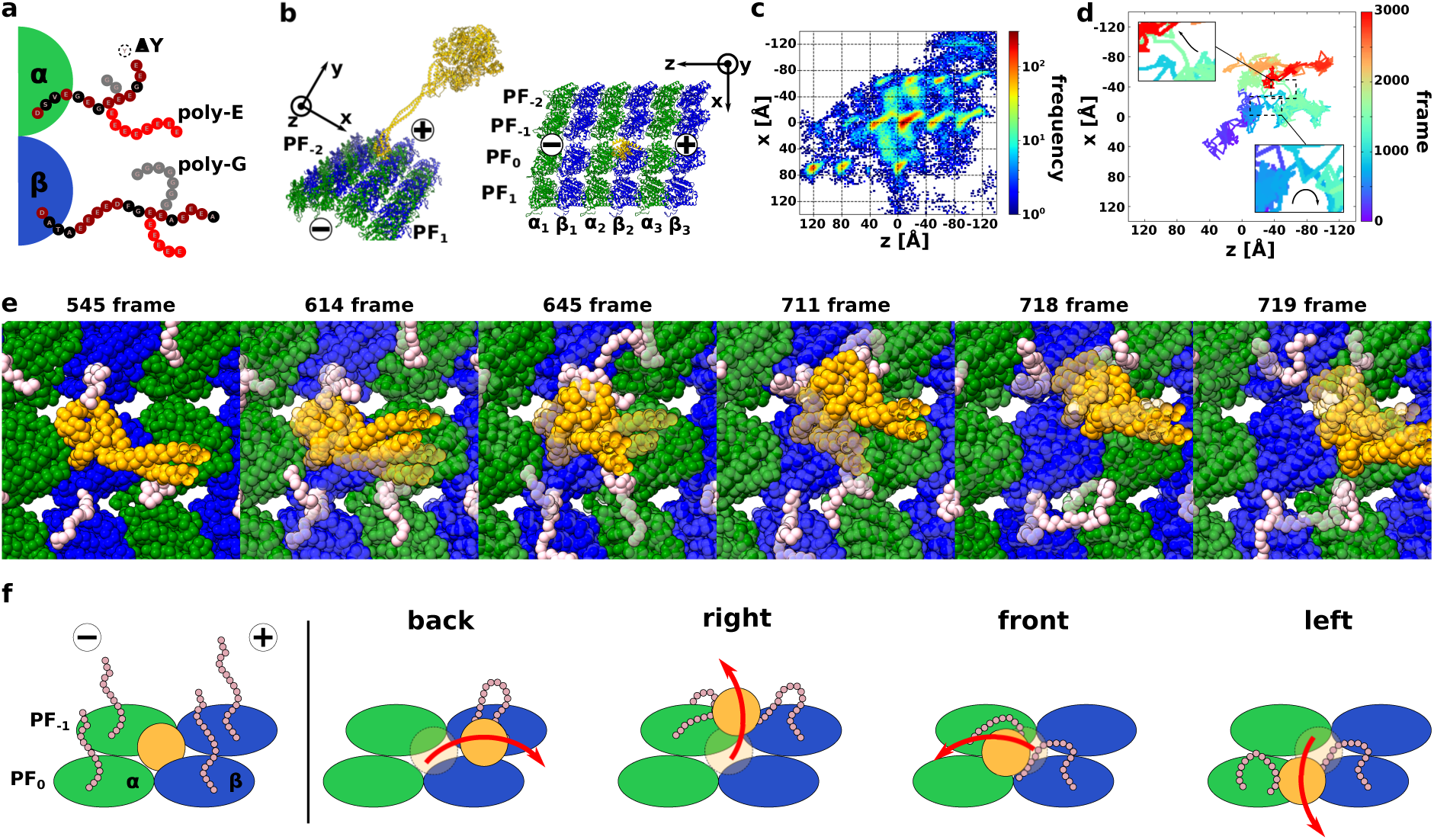
Low-affinity state dynein with WT MT. (a) PTM on CTT of tubulin. (b) Simulation system of dynein and MT. Dynein, α-tubulin, and β-tubulin colored orange, green, and blue. The minus end of the MT is along the z-axis, and the circumferential direction is x-axis. (c) Heat map of the position of the low-affinity state MTBD on the uMT, with 20 trajectories overlaid. The frequency increases from blue to red. The red area is the stable bound for MTBD, while the blue area is the unstable within the 20 times trajectories. (d) Representative trajectory of MTBD on the unmodified MT. The simulation time is 3000 frames, and the trajectory changed a color from purple to red from 0 to 3000 frame. Origin is the initial position of the center of mass of MTBD. (e) Snapshots of a back-stepping trajectory. These snapshots are picked up from Fig.1d bottom. CTT of α- and β-tubulin colored pink. The semi-transparent model is a previous snapshot. (f) Cartoon for the back, right-side, front, and left-side stepping mechanism.

A cilium is a membranated MT-based organelle, which has its own transport system, intraflagellar transport (IFT) for ciliary assembly and maintenance. IFT happens on ciliary doublet MT (DMT), which is composed of a 13-protofilament A-tubule and a partial 10-protofilament B-tubule (6). Due to space restriction within the cilia, IFT can only happen on less than 10 protofilaments in the outer region of the DMT. The DMT has very specific PTM signatures. The B-tubule is reported to be composed of detyrosinated, poly-glutamylated and mono- or poly-glycylated tubulins while the A-tubule has low level of PTM (7-11). It was observed that anterograde transport, powered by kinesin-2 moves along the B-tubule toward the tip of the cilia while the retrograde transport, pulled by dynein-2 towards the base of the cilia happens on the A-tubule (12). While PTM differences in A- and B-tubules are suggested as one of the reasons for anterograde and retrograde transport track exclusivity, there is still no concrete explanation for this phenomenon. It is unclear whether the track exclusivity is due to selective loading at the tips and base of the cilia or due to track preference controlled by the tubulin code or MAPs.

Molecular dynamic (MD) simulation can be used as a great tool for study the molecular mechanism of motor motility. Previously, the gold-standard atomistic MD simulations have been used for dynein in fluctuation analysis (13), and a study of the interactions with the MT (14). To simulate large scale dynamics, coarse-grained MD approaches have been successfully employed, such as inter-head coordination in dynein (15), conformational changing in dynein (16-18), stepping mechanism in kinesin (19, 20), and small globular proteins’ diffusive motion on MT with PTM (21). However, there has been not yet theoretical research for dynein stepping on MT with PTM.

In this work, we examined the motility of IFT dynein-2 on MTs to see whether PTM can play a role in IFT transport regulation. We performed MD of dynein-2 in high and low-affinity states moving on MTs with no-modification, detyrosination, poly-glutamylation and poly-glycylation. Our study reveals that the CTT and its PTM states plays a significant role in dynein directionality and interactions.

## Results

### Low-affinity dynein-2 shows side and back stepping

First, we set up a simulation system for dynein-2 motility on unmodified MT (also referred to as uMT). Our simulation system consisted of four rows of protofilament (PF_-2_ PF_-1_, PF_0_, and PF_1_) with three sets of α- and β-tubulin dimers (*α*_1_, *β*_1_, *α*_2_, *β*_2_, *α*_3_, *β*_3_), and a low-affinity state dynein in the center of the system (Fig. 1b, Materials & Methods). The initial MTBD position set above PF_0_-α_2_β_2_ in the center of the MT lattice (by using low-affinity state MTBD with tubulin complex: PDB ID: 3J1U, where almost close to the high-affinity MTBD bound position shown in PDB ID: 6KIQ). We performed 20 times independent MD simulations with this setup. Each trajectory has 3,000 frames with 1 frame equivalent to 10,000 MD steps. As the starting point for our simulation, the center of the MT lattice has the highest probability of MTBD localization (Fig. 1c). Due to the periodicity of the tubulin lattice, the entire MT is dotted with regions of high probability of localization similar to the center, namely “lattice like” probabilities. Our simulation shows that the low-affinity state dynein-2 moves not only along the initial PF_0_ but also to the left and right PFs (Fig. 1c, Fig. S1a). The probability of the presence of the MTBD on unmodified MT was (PF_-1_, PF_0_, PF_1_, outside lane) = (22%, 43%, 22%, 13%). To remove initial-status dependency, we didn’t use initial 500 frames in this calculation. Since the percentages of movement to the left and right PFs are almost the same, our result indicates that dynein-2 walked in a straight line overall while side-stepping left and right. This behavior is observed in *in vitro* single molecule assay with *Saccharomyces cerevisiae* dynein-1 (22, 23) and *Dictyostelium discoideum* dynein-1 (24). This is different from kinesins, which exhibit clear directional movement (25).

We looked further into the back-stepping mechanisms of dynein-2 (Fig. 1c). To know which residues are important for the back-stepping, we counted the number of contacts between CTT and MTBD, which are defined as residue pairs with an inter-residue distance of 1 nm or less. The CTT of PF_0_-α3, PF_-1_-α3, and PF_-1_-β2 showed the highest contact values with MTBD (Fig. S1b, c). From the contact map, it is expected that the PF_-1_-β2 CTT contacts MTBD first, carries MTBD backward, transfers MTBD to the adjacent PF_-1_- or PF_0_-α3. To clarify the detailed mechanism, we focused on the moment when the back-stepping occurs in a representative trajectory (Fig. 1d, e, Movie S1). First, after the MTBD moved 8Å backward (z=-8Å, x = 0Å), (Fig.1e, frame 545), the CTT of the PF_-1_-β2 attracted the MTBD (from frame 545 to frame 645). Subsequently, after contact occurred with the PF_-1_-α3 CTT (frame 711), the contact with PF_-1_-β2 was released (frame 718), and the 4 nm back-step was completed around frame 719. This trajectory details confirmed that the back-step was realized by the contact transition as expected.

Experimental data show that dynein-1 walks along MTs using a combination of coordinated and stochastic movement (26, 27) with a variable step size (23). Our simulation result indicates that the stepping behavior of dynein-2 is similar to dynein-1, which displays side and backward steps (23, 24). The backward step of chimeric yeast and *Dictyostelium* dynein-1 constitute of 20% and 14.7% of the total steps. Since the step-size and stepping probability are regulated by head-to-head separation (26, 27), our simulation with single dynein head represents only the stochastic nature of dynein stepping only without the influence of head-to-head coordination.

### The CTT-MTBD contact heavily influences the MT stepping direction

Since the CTT of tubulin is flexible and the CTT interaction with MTBD happens very quickly, these interactions are not measured by either single molecule or cryo-EM structure of dyneins and tubulins. MD simulation on the other hand allows the snapshot of interactions between the tubulin CTT and MTBD. The contact map of CTT residues with MTBD indicates clear differences among front, back, right and left stepping (Fig. S1b, Movie S2). For the right-side stepping, every tubulin on PF_-1_ showed highly contact with MTBD (Fig. S1b). Since the PF_-1_-β2 and PF_-1_-α3 showed high contact values in the back-step, it suggests a possibility that the initial movement of the right-side step is the same as that of the back-step. However, only in the right-side step, the PF_-1_-α2 has high contact (Fig. S1b). Therefore, the contact with the α-tubulin on PF_-1_ might be an essential factor for the right-side step (Fig. 1f). Using a representative trajectory, we observed that the center of MTBD mass positioned above PF_-1_ is essential for making this contact (Fig. S2A). The beginning is the contact with the β-tubulin CTT of PF_-1_, and then the timing of MTBD’s contact with the α-tubulin CTT will determine whether the right-side step or the back-step will be made.

The front and the left-side step show a high contact with the PF_0_-β2 which does not see in the back and right-side step (Fig. S1b). The front step also has a contact with the PF_-1_-α2, which is specific for the front step. Besides, the left-side step has a high contact with the PF_0_-α2, additionally (Fig. S1b). Therefore, it is expected that the front and left-side steps will exit the initial stable site by contacting the PF_0_-β2 at first, and then if the contact with the PF_-1_-α2 occurs, the front step will be occurred, and if the contact with the PF_0_-α2, the left-side step will be occurred. To confirm the actual molecular mechanism, we show the representative trajectory in Fig. S2b and snapshots in Fig. S2c, d.

Both the back-step and the right-side step are attracted by the CTT of the β-tubulin of the PF at the far end and comes to the edge of the area with a high probability of localization. The back-stepping occurs if MTBD contacts the CTT of the backward α-tubulin after MTBD reaches the end of the stable area (Fig. 1f, Fig. S3a). On the other hand, if this contact happens before MTBD reaches the end of the stable area, the right-side step occurs (Fig. 1f, Fig. S3b). For both the front step and the left-side step, MTBD needs to leave from a stable area at first. Then, if the β-tubulin CTT precariously sways MTBD at the far side, MTBD is positioned anteriorly and contacts the α-tubulin CTT at the front, and the front step occurs (Fig. 1f, Fig. S3c). If MTBD contacts with the α-tubulin CTT before it moves toward the front, the left-side step occurs by the contact with the α- and β-tubulin CTTs on the close side (Fig. 1f, Fig.S3d).

Our result indicates that the interaction of the CTT with the dynein-2 MTBD is critical for the stepping behaviors of dynein-2. *In vitro* data shows that chimeric yeast dynein-1 moves with higher speed with the removal of CTT from β-tubulin and both α- and β-tubulins (4). On the other hand, the CTT of tubulin is reported to increase processivity of dyneins and kinesins (4, 28). This agrees with our simulations that the tubulin CTT-dynein MTBD contacts heavily influence the dynein stepping direction and hence the run length and processivity. Therefore, the extensive contact of dynein-2 MTBD with tubulin CTT is a control mechanism for the motility of dynein-2.

### Detyrosination breaks the side-step ratio of low-affinity dynein

Next, we wanted to investigate how detyrosination changes the motion of dynein-2 in low-affinity state. We ran the simulation again after adjusting the MT in the setup to detyrosinated tubulins, referred to as ΔY MT (Fig. 2a, b). The heat map of MTBD locations clearly shows that the left and right side-step proportions are no longer equal compared to the unmodified MT (Fig. 2b). The frequency of dynein-2 on PF-_1_ increases 141% (31% vs. 22%) while the frequency of dynein on PF_1_ decreases 32% (7% vs. 22%) (Fig. 2b). These results indicate that detyrosination causes dynein-2 to move in a right-hand bias first stepping toward the minus end of MT.

**Figure 2.**
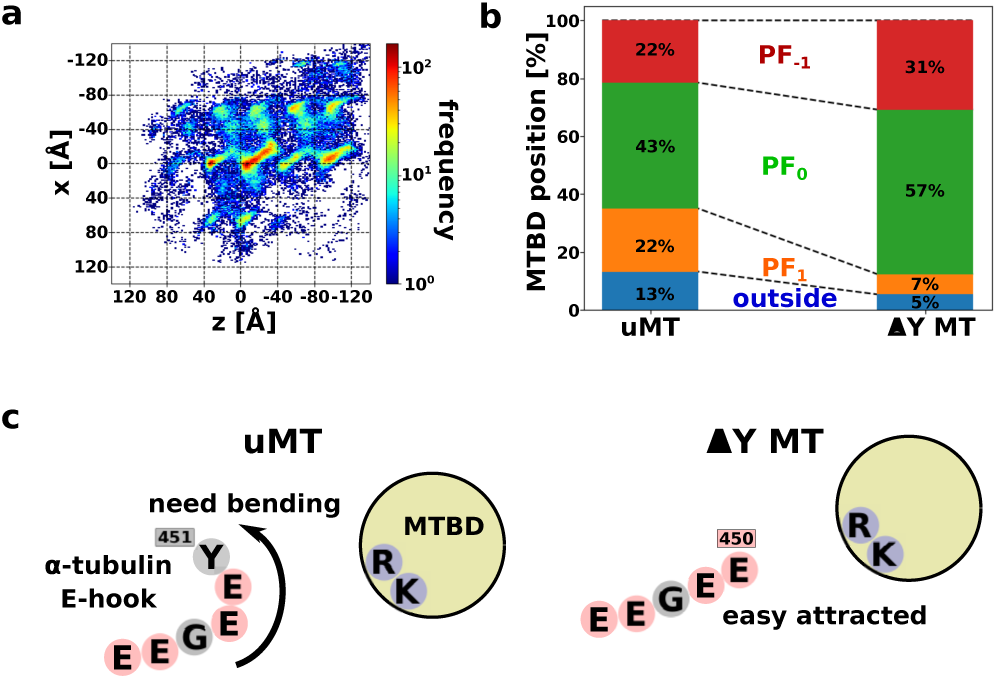
Low-affinity state dynein with detyrosinated MT. (a) Heat map of the position of the low-affinity state MTBD on the ΔY MT, with 20 trajectories overlaid. The coloring method is same with Fig. 1d. (b) Probability of localization of MTBD on the uMT and ΔY MT for each binding lane; PF_-1_, PF_0_, PF_1_, and other percentages are red, green, orange, and blue, respectively. (c) The detyrosinated CTT seems to have an easier contact MTBD than unmodified CTT. The positively charged ARG and LYS are blue, the negatively charged GLU and ASP are red, and the other are gray.

As shown in Fig.S1d, there was no clear difference in tubulin CTT – MTBD contacts between uMT and ΔY MT in the front-step. However, the contact with the CTT was increased for the other directional steps in ΔY MT. It is likely that the negatively charged E450, which was previously capped by Y451, is now exposed, making it more susceptible to the attractive effects of electrostatic interactions with positive charged arginine and lysine in dynein-2 MTBD (Fig. 2c, Fig. S1d). Since contact with α-tubulin is important for back-step and right-side step, the increase contacts due to detyrosination can lead to back-step and right-step biases.

Detyrosination is associated with robust motility of kinesin-2 *in vitro* and is shown to increase the velocity of kinesin (4). In the same study, chimeric yeast dynein-1 does not show different velocities on detyrosinated and tyrosinated MTs. *In vitro* data analysis shows that kinesin-2 exhibits left-stepping behavior on axonemal doublet MT in contrast to straight moving behavior of kinesin-1 (29). They implied that the stepping behavior of kinesin-2 is to avoid the retrograde cargo in the doublet MT and expected that dynein-2 also exhibits left-stepping bias for the same reason. Our simulation shows that detyrosination only influence the side-stepping ratio of dynein-2. In the doublet MT, the right-side step is likely not affected the runway significantly since it is not possible to step from A- to B-tubule. In addition, the behavior of human cytoplasmic dynein-2 on detyrosinated MT might be quite different from yeast. With a recent finding that detyrosination of tubulin does not affect the rate of MT assembly (30), it is thought that detyrosination is one way to fine tune the microtubule dynamics and interaction.

### Long poly-glutamate branch prevents MTBD from making contact with the globular domain of tubulin

Next, we examined the effect of polyglutamylation on dynein-2 motility. Since the number of glutamate residues attached to the CTT as branches varies from 1 to 6 in neuronal cells and as high as 10 or more in the ciliary doublet (10, 31-33), we used a wide range length of poly-glutamate (poly-E) branch to α- and β-tubulins for our simulations (Table 1).

**Table 1:** The combination of number of glutamate residues added to α- and β-tubulins in our simulation

| ID | 1 | 2 | 3 | 4 | 5 | 6 | 7 | 8 | 9 | 10 |
| --- | --- | --- | --- | --- | --- | --- | --- | --- | --- | --- |
| (α-, β-) | α-14E/β-18E | α-14E/β-0E | α-0E/β-18E | α-5E/β-0E | α-0E/β-5E | α-5E/β-5E | α-18E/β-18E | α-18E/β-5E | α-18E/β-0E | α-3E/β-18E |
| ID | 11 | 12 | 13 | 14 | 15 | 16 | 17 | 18 | 19 | 20 |
| (α-, β-) | α-3E/β-3E | α-3E/β-0E | α-18E/β-3E | α-0E/β-3E | α-5E/β-18E | α-8E/β-18E | α-8E/β-8E | α-8E/β-0E | α-18E/β-8E | α-0E/β-8E |

As shown in Fig.S4a, as the length of glutamate residues in both α- and β-tubulins increased, the lattice-like stepping became less and less visible. When the length of poly-E branch of either α- or β-tubulins was 18, the MTBD was almost completely retained at the starting position with no lattice-like motion (Fig S4a). This is because the MTBD is in a completely dissociated state from the MT.

When the length of poly-E branch is less than 8, we can barely see the lattice step (Fig. S4a, Fig. 3a) as shown in the unmodified MT (Fig. 1d), however, there are two new stable sites (boxes in Fig. 3a). To determine the locations of these new sites, snapshots were taken from a representative trajectory (Fig. 3b, c). One new stable position is on PF_0_ (Fig. 3c left, frame 540) with the CTT and the poly-E branch of these tubulins in contact with MTBD. This position is the intermediate state through which the left-side step is performed in the case of unmodified MT. While the contact with the PF_0_-β2 does not occur easily for the unmodified MT, the poly-E branch of PF_0_-β2 can reach out to MTBD. The second stable new site is on PF_-1_ of poly-E MT (Fig. 3c, frame 1516). It is the intermediate position on the right-side stepping in the case of unmodified MT. Like the first stable new site, poly-E branch can contact the α-tubulin side, which made it easier to go through the right-side step. However, due to the large number of contacts between the MTBD and the CTT of tubulins, it was not possible for MTBD to dissociate from flexible tails and therefore the new contact site was made.

**Figure 3.**
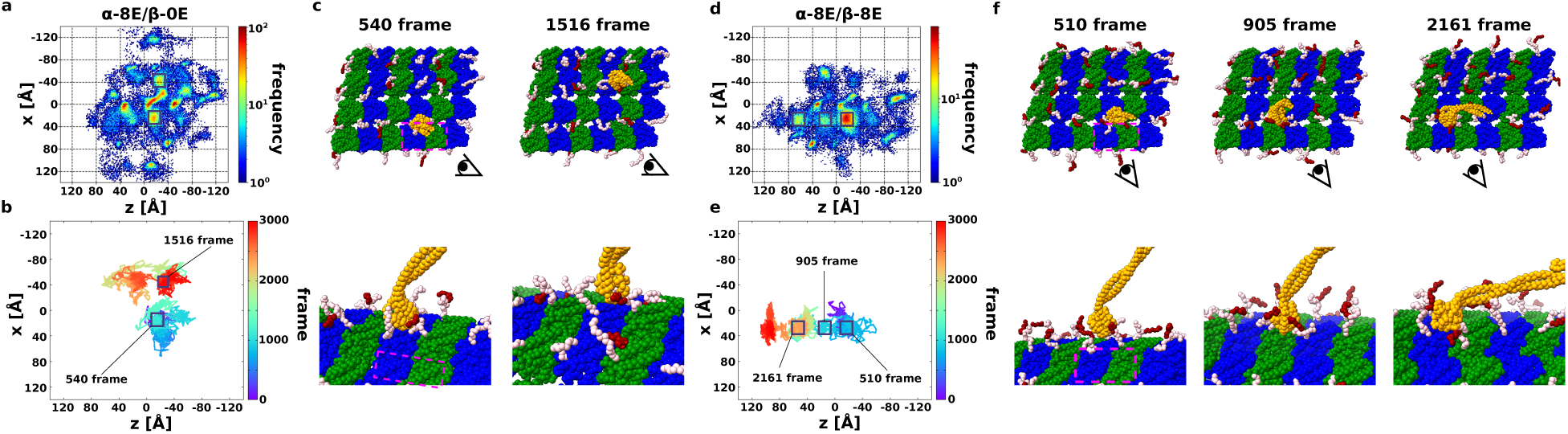
Low-affinity state dynein with short poly-E MT. (a, d) Heat map of the position of the low-affinity state MTBD on the poly-E MT, with 20 trajectories overlaid. The coloring method is same with Fig. 1d. (a) is α-tubulin have 8 length poly-E, and β-tubulin doesn’t have poly-E (simply denoted α-8E/β-0E). (d) is both α- and β-tubulin have 8 length poly-E (α-8E/β-8E). (b, e) Representative trajectory of MTBD on the poly-E MT. The coloring method is same with Fig.1b. (b) is the representative one in the case α-8E/β-0E, and (e) is the representative one in the case α-8E/β-8E. (c, f) Snapshots picked up from Fig.3b and Fig.3e inboxes. The coloring method of dynein and tubulins are same with Fig.1e, and poly-E colored red.

The motility inhibition becomes more pronounced as the length of the poly-E increases (Fig. S5). If there is no poly-E branch, only the CTT can freely contact MTBD. If only CTT can contact MTBD, MTBD can contact the globular domain of MT and perform lattice stepping. As the length of poly-E branch increases to 3, 5, 8, and 18, the number of contacts with CTT decreases, and by the time the length of poly-E branch is 8, the contacts with poly-E branch are dominant.

Interestingly, when comparing the effect of poly-E on α- and β-tubulins, we observed that the MTBD motility is affected more in the case of β-tubulin (Fig. S5). This can be because the CTT of β-tubulin is longer than that of α-tubulin, and the branching position of poly-E is located at the further C-terminal residues of β-tubulin than that of α-tubulin. Therefore, poly-E branch on β-tubulin can cover a wider radius. As a result, it is possible that the MTBD has completely dissociated from the MT. This dominant effect of β-tubulin over α-tubulin is also observed in the simulations of protein diffusion on MT with different tubulin codes (21)

In summary, by increasing the length of the poly-E branch for both α- and β-tubulins, the frequency of existing spots on MT decreased and the probability of the new spots increased. An important feature is that there are times when the MTBD is not in contact with the globular domain of tubulins and it is lifted by the poly-E branch and CTT (Fig. 3f, frame 510 and frame 2161). This is the reason why the density of lattices on MTs decreases as the poly-E length increases from 3 to 8 (Fig. S4a).

There is a lack of data on *in vitro* human dynein-1 and dynein-2 motility on PTM microtubules. Chimeric yeast dynein-1 shows no different on MT with 0, 3 and 10-E residues in both velocity and processivity (4). Our study shows that even short poly-E can affect the contact of MTBD to tubulin and significantly restricted contact in the case of long poly-E branch. Therefore, dynein-2 motility might be inhibited on poly-E MT. i.e., the B-tubule. It has been shown that hyper polyglutamylation can lead to defects in neuronal transport (34). This supports our results that long poly-E can inhibit motility significantly.

### Dynein-2 collapses onto MT surface in the case of long poly-E branch of β-tubulin

As shown previously that long poly-E branches restrict the movement of dynein-2, we wanted to investigate the different effects between the long poly-E of α- and β-tubulins. The heat maps and the representative trajectory of long poly-E of α-tubulin (α-18E/β-0E) and β-tubulin (α-0E/β-18E) cases show little differences with no step-like motion (Fig. 4ab and Fig. 4cd). However, the distance from the dynein-2 head to MT, namely head-to-MT is different. The head-to-MT distance did not change in the entire simulations in the case of unmodified MT (Fig. S4b) and the long poly-E on α-tubulin (Fig. 4e). In contrast, the head-to-MT gets shorter and shorter as time goes by distance in the case of the long poly-E on β-tubulin (Fig. 4f). The head-to-MT distance reduction mean that the entire dynein collapsed into the MT as shown in Fig.4f inset. This phenomenon was always observed when the poly-E branch of β-tubulin has 18 residues, regardless of the length of poly-E of α-tubulin (Fig. S4b). This suggests that while long poly-E for both α- and β-tubulins inhibits movement on MTs, long poly-E of β-tubulin may draw dynein-2 head to MT, so that it cannot properly bind to MT.

**Figure 4.**
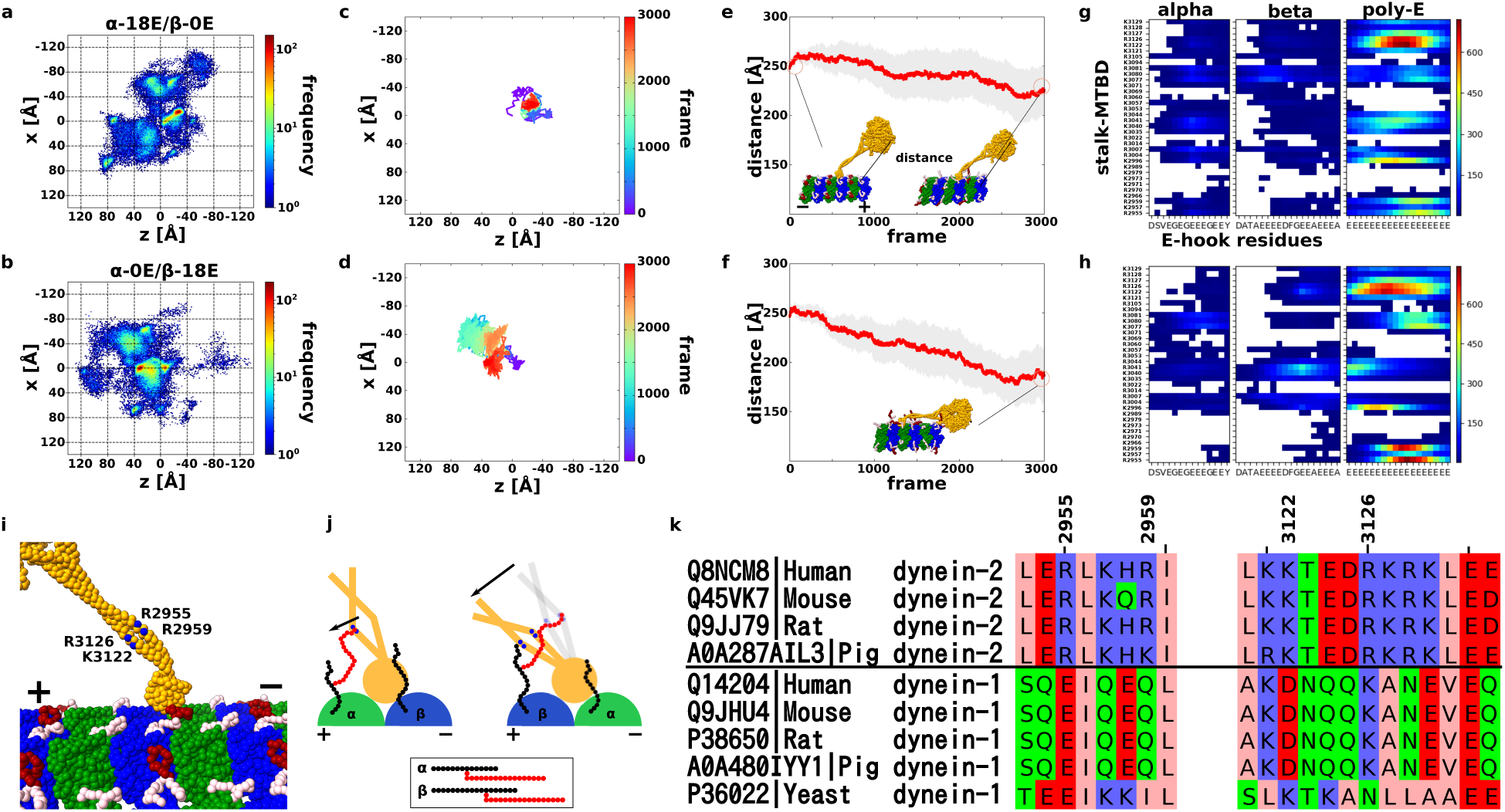
Low-affinity state dynein with long poly-E MT. (a, b) Heat map of the position of the low-affinity state MTBD on the long poly-E MT, with 20 trajectories overlaid. The coloring method is same with Fig. 1d. (a) is in the α-18E/β-0E case, and (b) is in the α-0E/β-18E case. (c, d) Representative trajectory of MTBD on the poly-E MT. The coloring method is same with Fig.1b. (c) is in the α-18E/β-0E case, and (d) is in the α-0E/β-18E case. (e, f) The trajectory of the average distance between the tip of the linker to the closest tubulins. The average distance for each frame is red, and 95% confidence interval are gray. (g, h) Residue-by-residue contact maps of CTT and poly-E with ARG or LYS residues on stalk-MTBD. Highly contacted pairs are colored red, lower are blue, and no contact pairs box are white. (g) and (h) is made from 20 trajectories in α-18E/β-0E and α-0E/β-18E setup, respectively. (i) The highest contact residues on the stalk-MTBD region; R2955, R2959, K3122, and R3126. Especially, R2955 and R2959 are specific contact residues in (0, 18) case. (j) Cartoon for understanding poly-E contact features. MTBD, α-tubulin, and β-tubulin colored orange (gray), green, and blue. CTT is black, and poly-E is red. (k) Sequence alignment of dynein-1 and -2 from different species.

What is the mechanism for this drastic phenomenon? To answer this question, we examined how α- and β-tubulin CTTs and poly-E branches contact with the arginine and lysine residues of dynein-2 stalk and MTBD. We observed many contacts with K3122 and R3126 and the poly-E tails for both α-18E/β-0E and α-0E/β-18E case (Fig.4g for α-18E/β-0E, and 4h for α-0E/β-18E, and 4i for both). In addition, R2955 and R2959 showed high contacts specifically for α-0E/β-18E case (Fig.4h).

Since the poly-E of β-tubulin can move in a wider radius than the poly-E of α-tubulin (Fig.4j). The poly-E of length 18 on the CTT of β-tubulin can contact not only the near side (K3122, R3126) but also the far side (R2955, R2959) of the coiled-coil of the stalk (Fig.4j right). On the other hand, a poly-E of length 18 on the CTT of α-tubulin can only contact the near side of the coiled-coil of the stalk (K3122, R3126) (Fig. 4j left). As a result, only the long poly-E branch of β-tubulin can draw the entire stalk and subsequently, the head to MT surface.

We also performed the simulation with dynein-2 MTDB in high-affinity state with poly-E MT (Supplementary Fig. 6, 7 and Supplementary Text). The high-affinity state dynein-2 is easily dissociating from MT without external forces or changes of nucleotide states in the presence of long poly-glutamylation. Therefore, long poly-E MT will impact the stable walking motility of dynein-2.

In the cilia, it is common to have more than 10 glutamate residues added to the CTT (10). We showed that long poly-E branch can pull dynein-2 stalk and subsequently head toward the MT. Therefore, the dynein-2 is likely to prefer the A-tubule to B-tubule due to lack of long poly-E branch in A-tubule. Interestingly, the residues important for the interaction with long poly-E branch (R2955, R2959, K3122, R3126) are conserved in dynein-2 from human, mouse, rat and pig but not dynein-1 (Fig. 4k). In addition, it seems like there are a cluster of four positively charged amino acids in the region of those important amino acids, which is not observed in dynein-1. This sequence difference suggests that dynein-1 and dynein-2 might exhibit different motility on poly-E MT. Therefore, the motility inhibition of long poly-E MT can be specific to dynein-2. In contrast to dynein, both the 3E- and 10E-CTTs increased the velocity and processivity of human and *C. elegans* kinesin-2 by 50%. Together, it suggests that kinesin-2 instead of dynein-2 is favoured on the heavily poly-glutamylated B-tubule.

### Poly-glycylation does not exhibit the same effect as poly-glutamylation

Next, we want to investigate how the motion of low-affinity state dynein changes on mono or poly-glycylated (poly-G) MT, a PTM specific to ciliary MT (35) (See setup in Materials & Method). In contrast to poly-E, there was no clear difference between unmodified MT and poly-G MT in the MTBD position heat map and the plot of the head-to-MT distance, suggesting that poly-G does not attract MTBD (Fig. 5a, b, Fig. S8).

**Figure 5.**
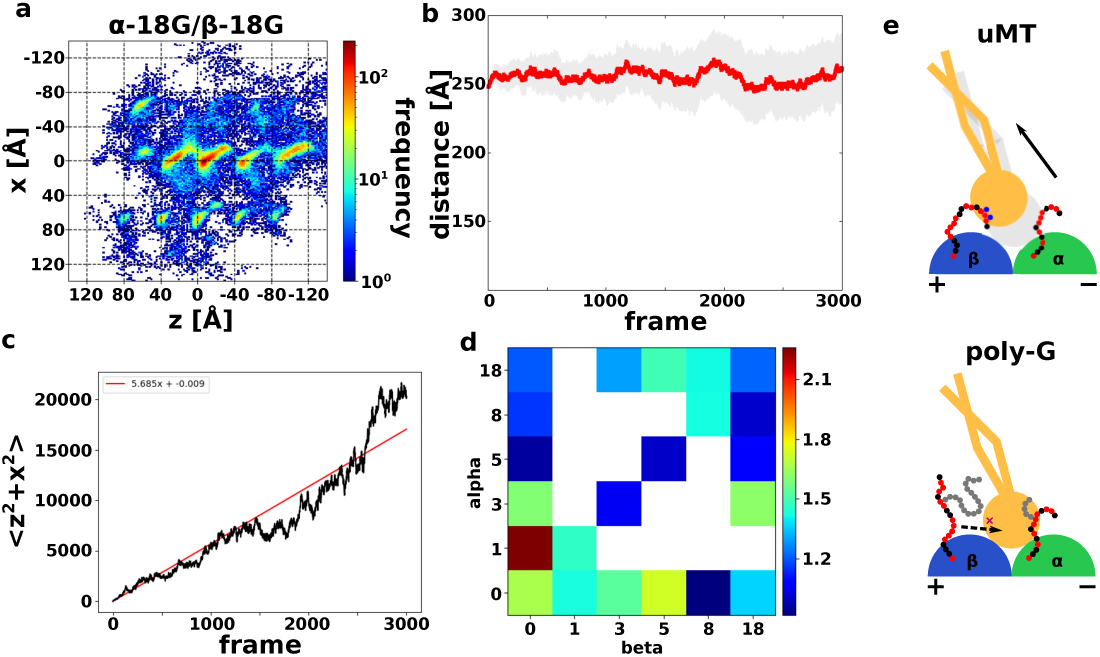
Low-affinity state dynein with poly-G MT. (a) Heat map of the position of the low-affinity state MTBD on the poly-G MT, with 20 trajectories overlaid. The coloring method is same with Fig. 1d. This is in the α-18G/β-18G case. (b) The trajectory of the average direction between the tip of the linker to the closest tubulins in the α-18G/β-18G case. The coloring method is same with Fig.4e. (c) MSD plot of α-18G/β-18G case (black) and its linear approximation line (red). (d) The heatmap of the diffusion coefficient in the various poly-G simulation systems. Blue is small diffusion, red is high diffusion, and the white is not simulated setups. (e) Cartoon for understanding poly-G simulation features. When MT doesn’t have poly-G, CTT directly contacts to MTBD, and CTT induces MTBD flexible motion (above). On the other hand, when poly-G MT prevents CTT from contacting MTBD, thus suppressing the diffusion of MTBD.

We predicted that the long poly-G might interfere the interactions between CTT and MTBD, making them less susceptible to the attractive effect of electrostatic interaction. If so, MTBD on the poly-G MT would be less likely to diffuse than in the unmodified MT because the main source of diffusion of low-affinity dynein is by CTT attraction. To investigate the degree of diffusion of MTBD, we made Mean Square Displacement (MSD) plots for each setup (Fig. S9). The diffusion coefficients were derived by linear approximation using the least squares method. Fig. 5c shows the MSD plot and linear approximation results for poly-G with a length of 8 for both α- and β-tubulins. The diffusion coefficients of each setup are shown as a heat map in Fig. 5d.

Poly-glycylation is shown to be essential for cell motility and division in *Tetrahymena thermophila* (36). However, that study showed that the total amount of poly-glycylation on both α- and β-tubulins is essential for survival instead of the poly-G amount on either α- or β-tubulins. Defects in glycylation in mouse sperm lead to aberrant motility of the cilia due to abnormal conformation of axonemal dyneins. However, since the length of the flagella in that mutant is almost the same as WT, IFT is unlikely affected. We suspect that the phenotypes of aberrant motility of axonemal dyneins in poly-G deficient mutant is likely due to binding of effectors in the cilia that affect axonemal dynein activity.

When doing simulation of the low-affinity state dynein-2 movement on the MT constructed with a combination of detyrosination, poly-E, and mono-G, resembling the human tubulin code (Fig. S10) (Janke & Magiera, 2020), poly-E displays as the dominant factor in impeding the stepping of the dynein-2 MTBD. This reinforces the notion that dynein-2 does not walk efficiently on the B-tubule which contains with detyrosinated and long poly-E tubulins.

## Discussion and conclusion

In this study, we showed that we can simulate the stochastic motility behaviors of dynein-2 on MT with different PTM. While we observed different behaviors in our simulation of a single head dynein-2, it is important to know there are any differences in the stepping mechanism between single-headed and double-headed dynein. Our results of the ratio of left- and right-side stepping is almost the same with the results of the experimental results of double headed dynein (24). Therefore, we expect no significant difference between single and double-headed dynein for the side-stepping. Since only one MTBD could contact each binding site on MT, if an MTBD is already bound in the direction where the other MTBD tries to step in the case of double-headed dynein, its movement will be hindered. In addition, when the head on the right side tries to take a left-side step, the left-sided head could get in the way and vice versa. However, in the low-affinity state dynein-2’s step, the MTBD might be moved by CTT or poly-E before the AAA+ ring region moves, so the side-step mechanism might not relate to the number of heads.

We predict front- and back-stepping mechanisms have a bit difference between single-headed and double-headed dynein. Since the lattice structure of MT does not change if dynein has a single head or not, it is assumed that there might be no significant difference in the locations that they pass through during the step. However, it is expected to be more difficult for the double-headed to move backward because the light chains are connected to the other head. To perform front-step in single-headed dynein, it was necessary to move to the energetically unstable forward site first. However, in double-headed case, when the other head performs the power-stroke motion, it is pulled forward through the tail domain, making it easier to clear the first step to perform the front step. This scheme is consistent with the experimental results showing that the linker’s power-stroke motion direction is essential for directional motion (37). In summary, the energetically stable backward step becomes more difficult with double-headed dynein, while the energetically unstable forward step becomes easier, which reverses the ratio of front-step to back-step, it is expected that the ratio of front-step will be dominant, and both side-step ratios will be equal, as observed in the experiment (24).

Our study concludes that dynein-2 does not favor to move on the B-tubule due to the heavily poly-E tubulins. Recently, it is reported that the IFT train moving toward the tip can turn to the base without reaching the ciliary tip when blocked by a physical barrier (38). Therefore, there is no need for an external factor to target the dynein-2 on to the A-tubule at the ciliary tip. The heavily polyglutamylated B-tubule can act as a mechanism to target dynein-2 and the retrograde IFT onto the A-tubule. In retrograde IFT, which moves on the A-tubule (12), the dynein-2 movement would be restricted in between PF A8 and A9 due to the presence of the outer dynein arms between PF A7/A8 and the outer junction build on PF A10. The lateral stepping of dynein-2 can only be restricted within these two PF lanes.

Single molecule study suggests that kinesin-2 is left-side bias to avoid collision with the IFT retrograde train (29). If the IFT anterograde moves on B-tubule, kinesin-2 can move on PF B1-B4. As a result, kinesin-2 can have more room for left stepping to avoid collision.

In summary, our study shows that we can study dynein-2 motility on MT with different PTMs using molecular dynamics. Our simulation system can also apply to other dynein species and gain insights into the molecular mechanism of general dynein motility, which can be tested experimentally.

## Materials & Methods

### Model building

We used the undecorated GDP MT structure resolved by cryo-EM for our MD simulation (39). Protein Data Bank ID was 6DPV. The sequence of α- and β-tubulin was set TUBA1B (UniProtKB: P68363) and TUBB (UniProtKB: P07437), respectively. We used clustal-omega (40, 41) for align these sequences, and we used MODELLER for making homology model (42).

In addition, we used human cytoplasmic dynein-2, of which sequence was taken from DYHC2 (UniProtKB: Q8NCM8). The reference structure of the ADP.Vi state dynein which took low-affinity MTBD was taken from the PDB ID: 4RH7 (43). This model was already contained whole motor domain, and the sequence was DYHC2, so we used MODELLER only for modeling disordered region. Also, for the ADP state dynein which took high-affinity MTBD was modeled by combining the three models taken from the PDB ID: 3VKH (44) for AAA+ and stalk, 3J1T (45) for stalk and MTBD, and 6KIQ (46) for MTBD and the interface with MT in the following protocol. At first, 3VKH was the X-ray crystal structure of *Dictyostelium discoideum* cytoplasmic dynein-1. While this model contained stalk and MTBD, its resolution was apparently low. On the other hand, 3J1T contained only stalk and MTBD region of *Mus musculus*. So, we integrated 3VKH with 3J1T via a molecular dynamics simulation as described (17). The model used in Kubo et al. (17) was already contained entire of the motor domain, but 3J1T was modelled by combined cryo-EM and molecular dynamics simulation. Recently, 6KIQ was reported by only cryo-EM, which was the complex of yeast cytoplasmic dynein MTBD at high-affinity binding state with tubulin. Therefore, we integrated the model we already made with 6KIQ via Coot. We used clustal-omega for align these sequences to DYHC2, and we used MODELLER for making homology model (42).

We also modelled the three posttranslational modifications of tubulins: (1) detyrosination (delta-Y) of α-tubulin. (2) poly-glutamylation (poly-E) of α- and β-tubulins. (3) poly-glycylation (poly-G) of α- and β-tubulin. At first, delta-Y only eliminated the tyrosine at the C-terminal end of α-tubulin. Secondly, poly-E was modeled by MODELLER as a series of 18 glutamic acids, and then shortened as necessary. The N-terminal end of poly-E was placed at 3.6 Å from E445 of α-tubulin and E438 of β-tubulin. These positions of the branches were determined from (47) and bovine (48) for α- and β-tubulin, respectively. Lastly, poly-G was modeled by the similar procedure as poly-E. The N-terminal end of poly-G was placed at 3.6 Å from E446 of α-tubulin and E439 of β-tubulin. These positions of the branches were next to the position where poly-E branch started (49). Due to the absence of functional TTLL10 in human, poly-G should take mono-G (50), but we simulated not only mono-G but also poly-G for controlling.

### MD simulation for low-affinity dynein

We simulated the dynein motion on MT with and without PTMs. Fig 1b shows the overall setup of our simulation system: We set four PFs containing three α--β-tubulin dimers, and one dynein. The initial dynein was placed on PF_0_-αβ2 based on PDB ID: 3J1U. Table 1 shows the setting of the simulated PTMs. As shown in the Table 1, poly-E simulation was set to 0, 3, 5, 8, 14 and 18 length cases for α- and β-tubulin respectively. Similarly, poly-G simulation was set for lengths of 0, 1, 3, 5, 8, and 18. We also performed delta-Y simulation by removal C-terminal end of α-tubulin tyrosine as mentioned above. In each simulation, we performed 20 MD runs using CafeMol version 2.1 (51). Unless otherwise noted, we took 3 × 10^7^ MD steps. We used the underdamped Langevin dynamics at 323K temperature. We set the friction coefficient to 2.0 (CafeMol unit), and default values were used for others.

### Step definition

We defined four types stepping motion; front-, back-, left-side, and right-side step, for picking up each stepping from trajectories. At first, we defined the back-step as an intermediate state when the MTBD is located at x is in (−30, 30) and z is in (−38, -42) (Fig.S1a). That is because we can see that z = -40 is clearly the boundary between the initial stable position and the back-side stable position (Fig.1c). Next, we defined the right-side step from PF_0_ to PF_-1_. We defined this right-side step as an intermediate state when the MTBD is located at x is in (−28, 32) and z is in (−120, 120) (Fig.S1a). That is because we can see that x = -30 is clearly the boundary between the initial stable position and the right-side stable position (Fig.1c). Same as before, we defined the front-step and left-side step (PF_0_ to PF_1_) as an intermediate state when the MTBD is located at x is in (−30, 30) and z is in (−2, 2), and x is in (28, 32) and z is in (−120, 120), respectively (Fig.S1a). Those are because we can see that z = 0 and x = 30 is clearly the boundary for each stepping between the front-/left-side step and initial stable position, respectively (Fig.1c).

## Data Availability

The cryo-EM structures used in this paper are available download from the Protein Data Bank under PDB IDs 4RH7 for the low affinity state dynein; 3VKH, 3J1T, 6KIQ for the high affinity state dynein; 3J1U for the initial position definition of low affinity state dynein; 6DPV for uMT.

All MD simulations in this paper was performed by CafeMol software. It can be downloaded from https://www.cafemol.org.

## Author Contributions

SK and KHB conceived the project. SK designed, performed the MD simulation, analyzed the data, made figures and write the manuscript.

## Acknowledgements

We thank Drs. Muneyoshi Ichikawa, Shoji Takada and Anthony Roberts for critically reading the manuscript. SK is supported by JSPS KAKENHI Grant Number JP21J00021. KHB is supported by the grants from Canadian Institutes of Health Research (PJT-156354), Natural Sciences and Engineering Research Council of Canada (RGPIN-2016-04954).

