## Supplementary Text and Figures for "Regulatory mechanisms of the dynein-2 motility by post-translational modification revealed by MD simulation"

### Supplementary Information

#### Supplementary Text

##### The atomic interaction-based coarse-grained (AICG2+) model

We performed a coarse-grained MD simulation using the model mentioned above. We used a previously-developed and well-tested coarse-grained model AICG2+ (1, 2), in which each amino acid was represented by a single bead located at their Cα position. In this model, the reference structure was treated as the most stable conformation, and their parameters inside were defined from the all-atom reference structures. The energy function was written as

$$V_{AICG2+}(R|R_0) = \sum_i K_{b,i}\left(b_i - b_{i,0}\right)^2 + V_{loc}^{flp} + \sum_{j=i+2} \varepsilon_{loc,ij} exp\left(-\frac{\left(r_{ij} - r_{ij0}\right)^2}{2W_{ij}^2}\right)$$
$$+ \sum_{j=i+3} \varepsilon_{loc,ij} exp\left(-\frac{\left(\phi_{ij} - \phi_{ij0}\right)^2}{2W_{\phi,ij}^2}\right) + \sum_{i<j-3}^{nat\ contact} \varepsilon_{go,ij}\left[5\left(\frac{r_{ij0}}{r_{ij}}\right)^{12} - 6\left(\frac{r_{ij0}}{r_{ij}}\right)^{10}\right]$$
$$+ \sum_{i<j-3}^{non-native} \varepsilon_{ev}\left(\frac{d}{r_{ij}}\right)^{12}$$

In the order, each term represented the elasticity of the virtual bond between two consecutive Cα's, the sequence-dependent local potential made of virtual-angle- and virtual-dihedral-angle terms, the structure-based local potential between i-th and i+2-th residues, the structure-based local potential for dihedral angles, the contact potential for non-local natively interacting pairs (called "Go potential"), and the generic repulsion for the rest of non-local pairs. The vector *R* stood for $3n_{aa}$-dimensional Cartesian coordinates of the target protein where $n_{aa}$ was the number of amino acids in the protein. $R_0$ was the corresponding coordinate at the reference structure (All the variables with the subscript 0 meant the parameters that took the corresponding values at the reference structure). $b_i$ was the i-th virtual bond length between i-th and i+1-th amino acids. $r_{ij}$ was the distance between the i-th and j-th residues. $\phi_{ij}$ was the dihedral angle defined as i-th, i+1-th, i+2-th, and i+3-th residues. $W_{\phi,ij}^2$ was the parameter representing the width of the attraction. $K_{b,ibd}$, $\varepsilon_{loc,ij}$, $\varepsilon_{go,ij}$ were parameters which evaluated by AMBER force field via a multiscale algorithm. $\varepsilon_{ev}$ and $d$ were determined from a structural survey. The default values of these parameters and the meaning of them were written in the CafeMol manual (3).

For the disordered region, there were no reference conformation. To describe this feature, we set the Go potential in the disordered region to zero. By setting it zero, the disordered region behaved according to the $V_{loc}^{flp}$ term. This term was called flexible local potential, which was the sequence-dependent local potential (4). By using this function, poly-E and poly-G regions, which do not have the reference structure, would behave differently according to their respective sequences.

#### Parameter determination for high-affinity dynein

The stall force of dynein was already measured by single molecule experiments. There was no data for human dynein-2, but mammal dynein would detach from MT when dynein pulled toward minus end of MT under 1-3 pN (5, 6). Also, many single molecule experiments shown that dynein had asymmetric detachment rate; dynein easily detach when it is pulled toward minus end of MT and it hardly detach when it is pulled toward plus end of MT (7). So, in our coarse-grained MD simulation, high-affinity state dynein will detach when 1-3 pN force applied toward minus end of MT, and also needed to show the asymmetric detachment rate.

For controlled attractive force between MTBD and MT, which demonstrates the asymmetric detachment rate, we adjusted the Go potential. For adjustment, we prepared two PF rows of one αβ-tubulin dimer, and placed dynein in the center of the rows. Since the PDB structure 6KIQ is a complex model of an α--β- tubulin dimer with MTBD, dynein was placed based on the conformation in 6KIQ. We prepared four different Go potential scales: 0.2, 0.18, 0.15, and 0.1 times the default value. Additionally, for each Go potential scales, we prepared nine scale external force: -4, -3, -2, -1, 0, 1, 2, 3, and 4 pN. Minus value meant the force is applied to the plus end of MT. Finally, we performed 30 MD runs using CafeMol version 2.1 (3). We took $3 \times 10^7$ MD steps for each Go potential values and for each pulling force scale. All other setups were similar to the setup in the case of low-affinity dynein simulation.

When we used default Go potential values, dynein couldn't detach from MT (data not shown). When we used small Go potential, 0.15- or 0.1-times default value, we could not get asymmetric detachment rate (Fig. S6). However, when we used 0.2- or 0.18-times default value for the setting Go potential, we could get asymmetric detachment rate and detachment of dynein started around

2 pN (Fig. S6). Therefore, we determined 0.2 times default value was the best parameter for attractive force between high-affinity dynein with MT.

#### MD simulation for high-affinity dynein

To observe how the behavior of dynein on MT changes under the influence of PTMs when dynein in the high-affinity state was not bound to the strong binding site, the initial position of dynein was moved 1 nm above the strong binding site, which was placed on $PF_0$-αβ2 based on PDB ID: 6KIQ. To observe the effect of PTM when dynein in the high-affinity state was dragged by an external force, we made simulations where no force and a force of 3 pN was applied in each of the plus and minus end directions of MT. Then, we performed 10 MD runs using CafeMol version 2.1 (3). We took $3 \times 10^7$ MD steps for each pulling force and for each PTM conditions. All other setups were similar to the low-affinity dynein simulation case.

#### High-affinity dynein-2 easily dissociates from poly-E MT

Dynein needs to control two different structural states with different affinities to MTs to walk on them. Our earlier simulations showed that dynein MTBD in low-affinity state is affected by PTM. We now wondered whether dynein MTBD in a high-affinity state is affected by PTM. We first examined the contact force between MTBD and MT. Although the complex structure of dynein and tubulins in the strongly bound state has already been obtained (PDB ID: 6KIQ), it is necessary to confirm whether the degree of binding can be reproduced correctly in the coarse-grained simulation. In this study, we focused on the direction-specific dissociation of dynein in the strongly bound state. It is known that dynein in the strongly bonded state does not dissociate easily when subjected to a force in the plus end direction but dissociates easily when subjected to a force in the minus end direction. Since no direction-specific dissociation constants or stall forces have been measured in human dynein-2, we assume that the dissociation occurs with a force of about 2 pN in the minus end direction (5). By adjusting the contact strength (called coef-Go) between the MTBD and the tubulins, we were able to confirm that the dynein dissociated to the minus end but not to the plus end by applying an external force of about 2 pN when we set coef-Go to 0.2 times the default value (Fig. S6). The details were described in Method.

Based on the obtained contact forces between the MTBD and tubulins, we simulated how dynein in the strongly bound state behaves on MTs with PTMs. First, to improve the efficiency of the simulations, the entire dynein-2 model was moved 1 nm in the y-axis direction (away from the MT surface) from the initial binding position obtained by PDB model. Using this state as the initial structure, we performed 10 simulations for each setup. To investigate the effect of poly-E on α- and β-tubulin, we prepared eight different systems with poly-E on either side of α- and β-tubulin: α-0E/β-3E, α-0E/β-5E, α-0E/β-8E, α-0E/β-18E, α-3E/β-0E, α-5E/β-0E, α-8E/β-0E, and α-18E/β-0E.

Dynein-2 on unmodified MT immediately recontacted with MT and hardly moved from the initial position (Fig. S7ab). On the other hand, the longer the poly-E of MT, the more trajectories were observed to move from the initial position (Fig. S7cde). Moreover, we found the dissociation was triggered by contact with poly-E even after MTBD bound to the MT. As a representative example, the trajectory and heat map of the MTBD position in unmodified MT and α-0E/β-18E are shown in Fig. S7abcd. When snapshots were taken before and after dissociation from Fig. S7d trajectory, poly-E was entangled in the MTBD and pulled apart as shown in Fig. S7f. Overall, there is more movement in the case of β-tubulin poly-E than α-tubulin poly-E.

These results suggested that the high-affinity state dynein-2 is easily detached from MT without external forces or changes of nucleotide states in the presence of long poly-glutamylation. To achieve bipedal locomotion, one head must be weakly bound to the MT while the other head is strongly bound. However, in the presence of poly-E, even if the one head is bound in the strong binding state, that head is easily dissociated from the MT, as shown in Fig. S7g, and so, the stable walking motion cannot be realized.

Our results of high-affinity dynein-2 in the case of poly-E both reinforce the notion that dynein-2 does not walk efficiently on MT with long poly-E. While our simulation can only perform with single-headed dynein-2, we expect that double headed dynein-2 motility on the poly-E MT will not be much different due to the interaction of the MTBD with long poly-E branches.

#### Long poly-E is the dominant PTM for dynein-2 motility

Finally, we simulated the low-affinity state dynein-2 movement on the MT constructed with the human tubulin code. The human tubulin code consisted of detyrosination, poly-E, and mono-G (Fig. S10a) (8). Since only the length of poly-E branch was a variable site in this system, we performed 20 times simulations for each of 3, 5, 8, and 18 poly-E for both α- and β-tubulins (Fig. S10b and c).

When poly-E branch was short (α-3E/β-3E), the percentage of left-side stepping (presence on $PF_1$) slightly increased compared to ΔY MT (Fig. S10c). This is because the newly generated position with high frequency in the presence of poly-E, discussed in Fig. 3, was created on the $PF_1$ side. In fact, α-3E/β-3E and α-3E/β-0E in Fig. S4a, we can see the high-frequency position closer to $PF_1$, which was the same as created by α-3E/β-3E in Fig. S10b. However, this change was not enough to significantly affect the dominant feature of the right side-step toward $PF_{-1}$ by ΔY MT. The probability of the localization of right side-stepping ($PF_{-1}$) is still higher than left-side stepping ($PF_1$). Compared to ΔY MT itself shown in Fig. 2b, the frequency of lattice-like structures due to the shape of MT is less visible. Therefore, short poly-E or mono-G might make dynein-2 hardly bound to MT compared with unmodified MT and ΔY MT.

When poly-E became a little longer (α-5E/β-5E and α-8E/β-8E of Fig. S10b), the probability of left-side stepping ($PF_1$) became larger and larger (Fig. S10c). In the case of α-5E/β-5E, we can see the lattice-shaped high probability position in $PF_0$. In addition, the percentage of the outside from MT was not much different from that of ΔY MT or α-3E/β-3E. Therefore, in the case of α-5E/β-5E, poly-E, CTT, and the globular region of MT kept binding with dynein-2, and the diffusive motion was observed. However, in the case of α-8E/β-8E, the frequency of lattice formation due to the shape of the MT was no longer visible, and the rate of outward protrusion was drastically reduced. This means that the dynein-2 got strong contact with poly-E, CTT, and MT, so the diffusive motion of dynein-2 was reduced.

Finally, when poly-E was long (α-18E/β-18E), the percentage of left-side stepping ($PF_1$) decreased and right-side stepping ($PF_{-1}$) increased (Fig. S10c). The MTBD no longer stays in a stable position with a high frequency. The lattice-like frequency is no longer observed, which means that dynein-2 is no longer in contact with the globular domain of MT and is only trapped in poly-E and CTT. Since there is no contact with the globular domain of MT, dynein-2 cannot stay stably at the new binding sites on MT. Moreover, there are no characteristic high-frequency sites since dynein-2 drifts away from MT due to the flexible poly-E and CTT.

As in poly-E, the collapse of dynein-2 into MT was observed when poly-E was long (Fig. S10d). These results indicate that the low-affinity state dynein-2 on the human tubulin code MT exhibited similar dynamics as observed in the system with poly-E alone if the poly-E is long. Therefore, long poly-E is the dominant factor in the case of dynein-2 motility. This reinforces the notion that dynein-2 does not walk efficiently on the B-tubule which contains with detyrosinated and long poly-E tubulins. This supports the hypothesis that retrograde IFT transport by dynein-2 is on A-tubule, where there is mostly no modification.

### Supplementary Figures

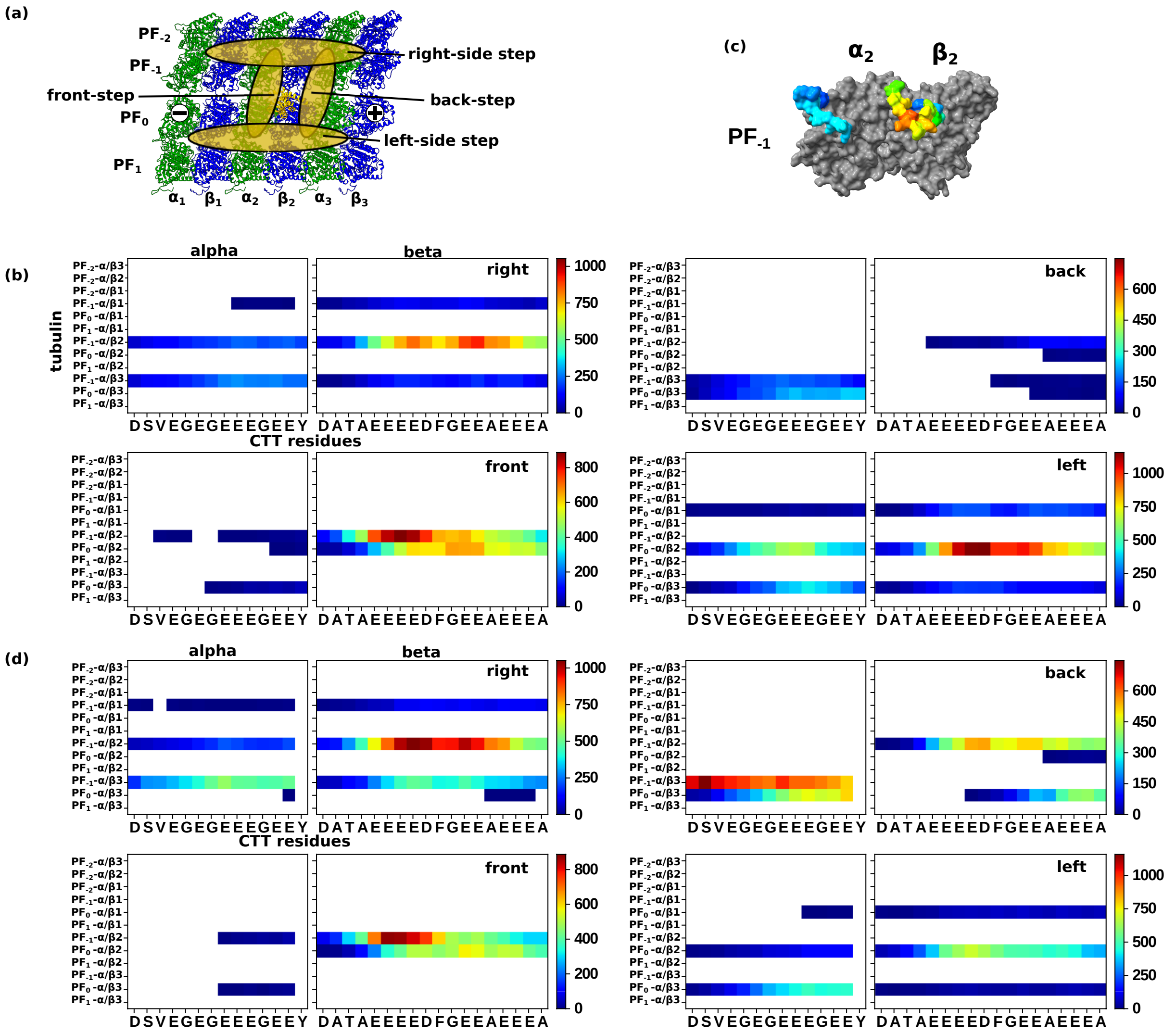


**Figure S1 Low-affinity state dynein with uMT and ΔY MT.**

(a) The definition area for four types stepping. (b, d) Contact map between CTT of α/β-tubulin and MTBD for each position; right-side step, back-step, front-step, and left-side step. Highly contacted region colored red, and lower is blue, and no contacted region is white. (b) In the case of uMT. (c) Mapping $PF_{-1}$ α/β2 of uMT right stepping contact value to tubulins for better visualization (d) In the case of ΔY MT.

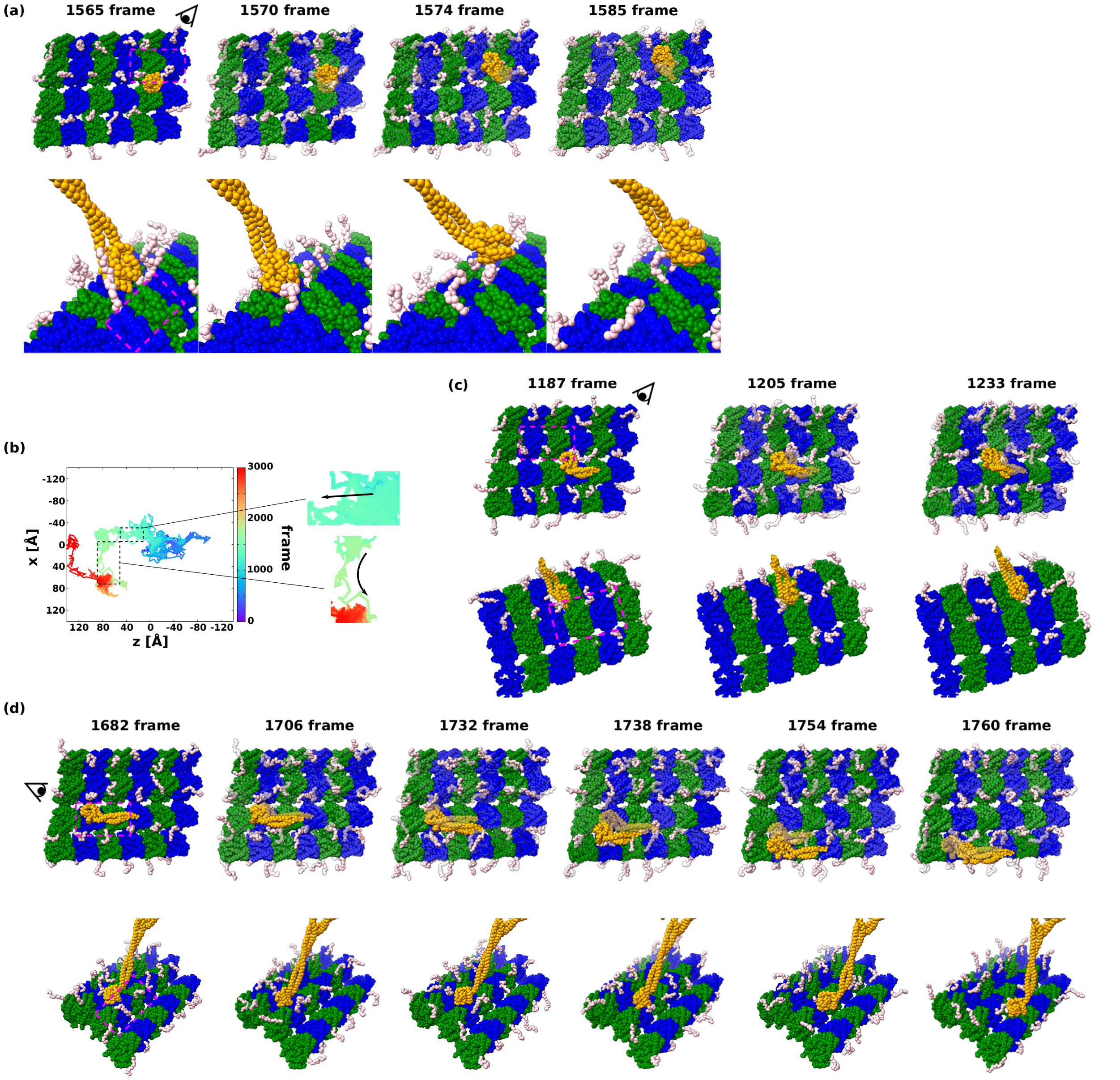


**Figure S2 Snapshots of representative stepping on unmodified MT.**
(a) Snapshots for the right-side stepping. These snapshots are picked up from Fig.1d above. (b) The other representative trajectory for picking up front-stepping (inbox above) and left-side stepping (inbox bottom). (c, d) Snapshots for the front-stepping and left-side stepping picked up from Fig.S2b.

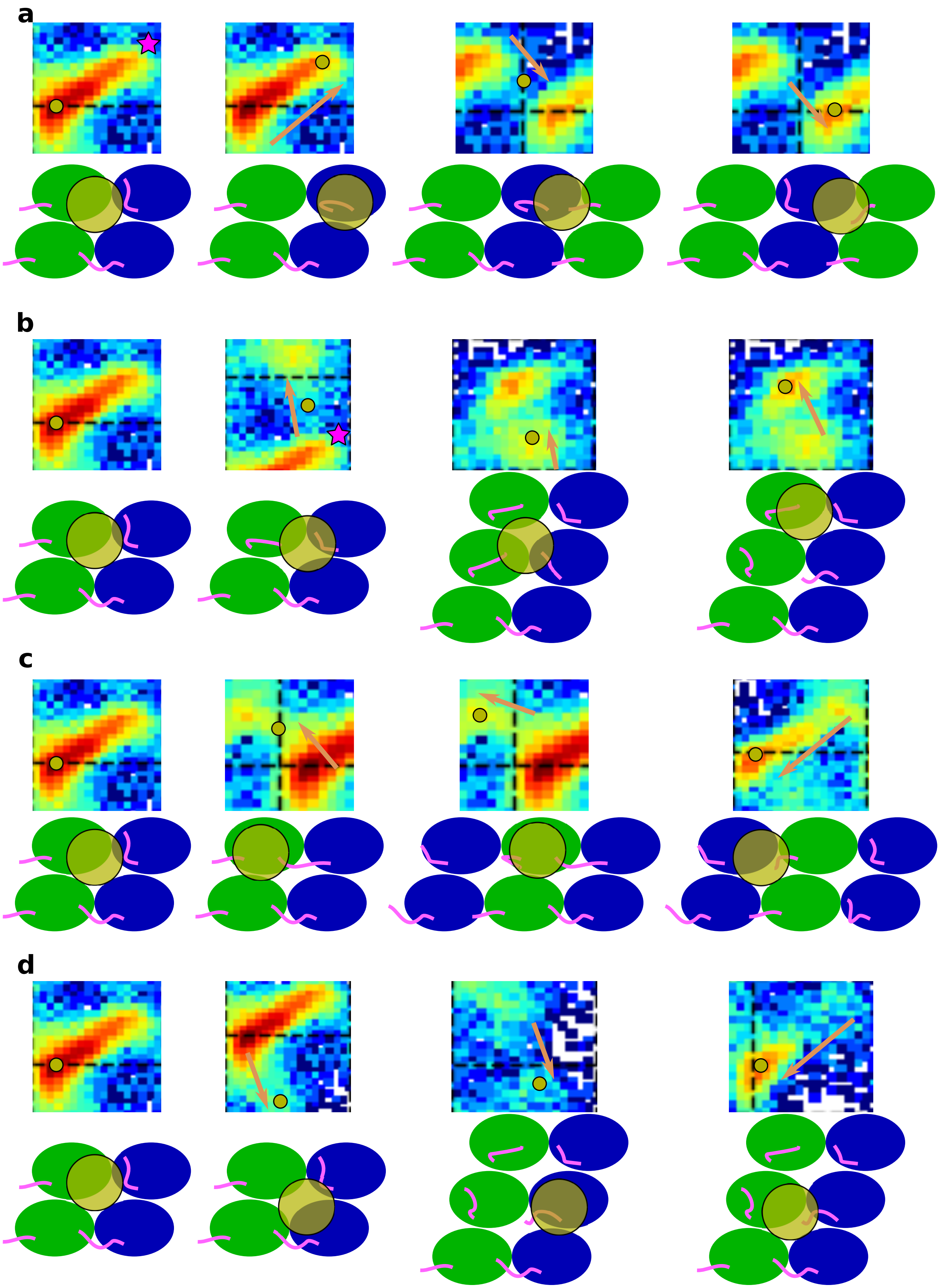

**Figure S3 Cartoon for four types stepping motion on the unmodified MT.**
Picking up focused region from the MTBD position heatmaps and the cartoons of specific molecular mechanism. (a) back-step (b) right-side step (c) front-step (d) left-side step. The magenta star in (a) and (b) is the end of the stable area.

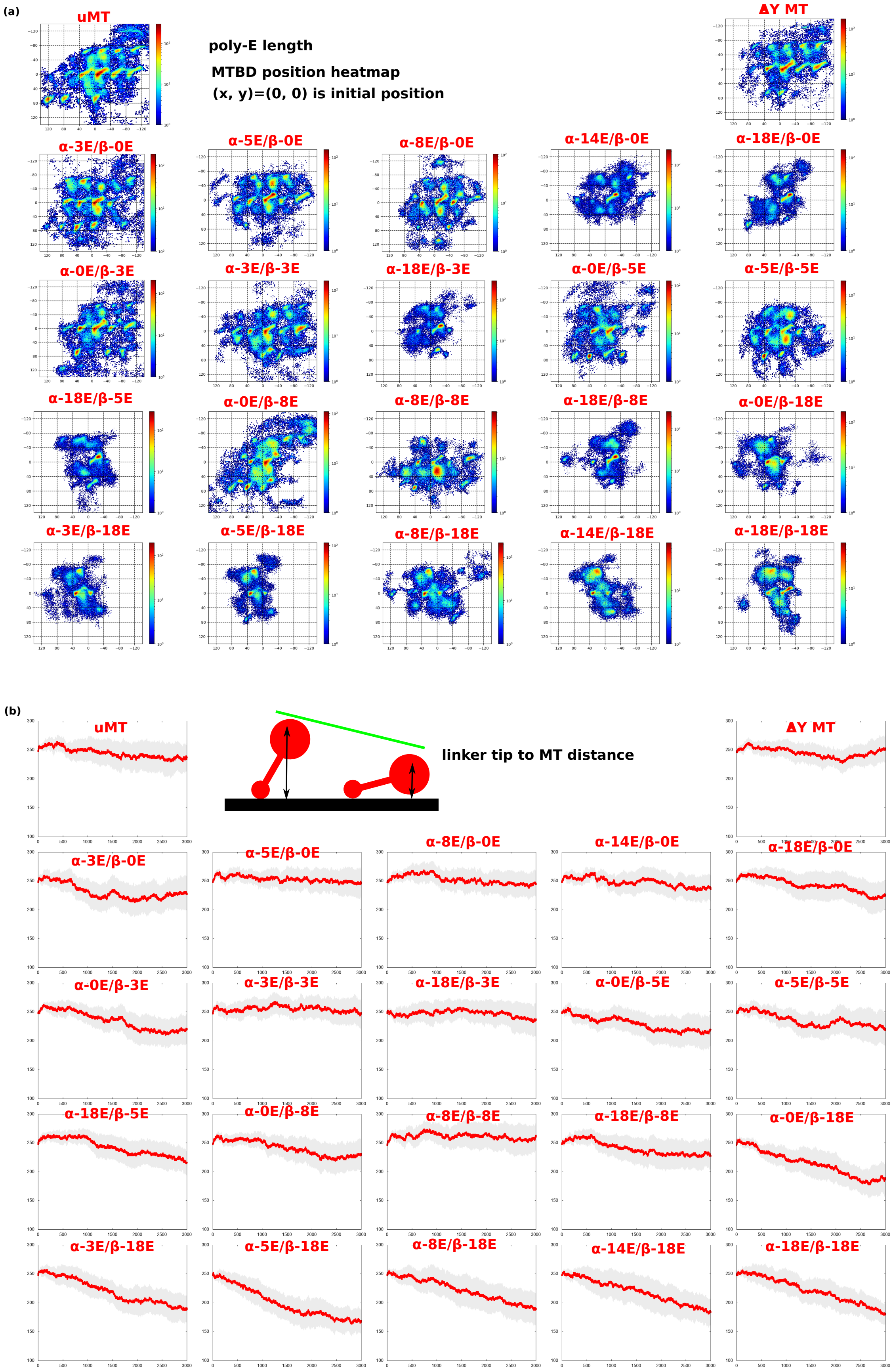
(a)
uMT
ΔY MT
poly-E length
MTBD position heatmap
(x, y)=(0, 0) is initial position
α-3E/β-0E
α-5E/β-0E
α-8E/β-0E
α-14E/β-0E
α-18E/β-0E
α-0E/β-3E
α-3E/β-3E
α-18E/β-3E
α-0E/β-5E
α-5E/β-5E
α-18E/β-5E
α-0E/β-8E
α-8E/β-8E
α-18E/β-8E
α-0E/β-18E
α-3E/β-18E
α-5E/β-18E
α-8E/β-18E
α-14E/β-18E
α-18E/β-18E
(b)
uMT
linker tip to MT distance
ΔY MT
α-3E/β-0E
α-5E/β-0E
α-8E/β-0E
α-14E/β-0E
α-18E/β-0E
α-0E/β-3E
α-3E/β-3E
α-18E/β-3E
α-0E/β-5E
α-5E/β-5E
α-18E/β-5E
α-0E/β-8E
α-8E/β-8E
α-18E/β-8E
α-0E/β-18E
α-3E/β-18E
α-5E/β-18E
α-8E/β-18E
α-14E/β-18E
α-18E/β-18E

**Figure S4 Low-affinity state dynein MTBD position heatmap and direction between the tip of linker and MT.**
(a) MTBD position heatmap for each setup. (b) Average distance between the tip of linker and the closest tubulin for each setup. Red line is the average of 20 trajectories for each frame, and gray is 95% confidence interval.

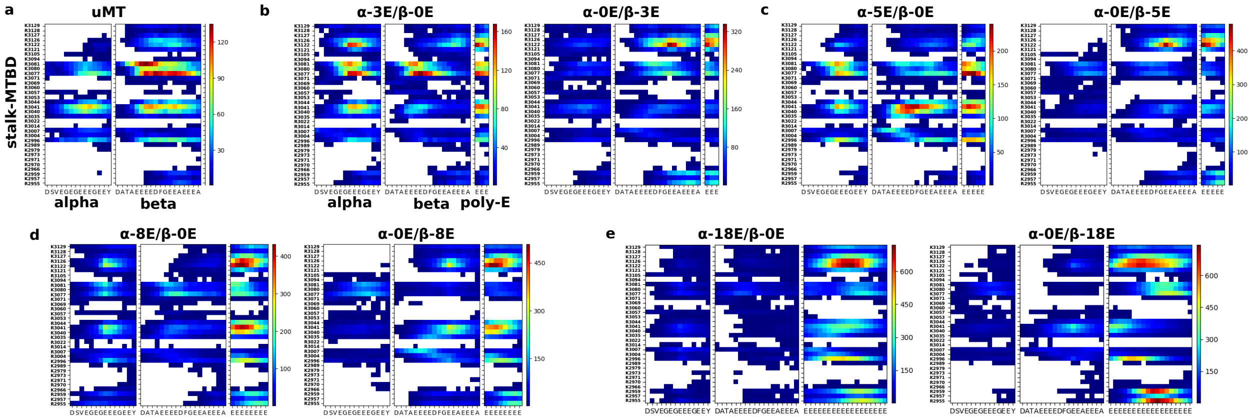


**Figure S5 Contact map between CTT/poly-E and stalk-MTBD.**
Contact map between CTT/poly-E of α/β-tubulin and stalk-MTBD for each poly-E setup. Highly contacted region colored red, and lower is blue, and no contacted region is white. (a) In the case of uMT. (b) In the case of 3 length poly-E MT. Left panel is α-3E/β-0E, and the right panel is α-0E/β-3E. (c) In the case of 5 length poly-E MT. Left panel is α-5E/β-0E, and the right panel is α-0E/β-5E. (d) In the case of 8 length poly-E MT. Left panel is α-8E/β-0E, and the right panel is α-0E/β-8E. (e) In the case of 18 length poly-E MT. Left panel is α-18E/β-0E, and the right panel is α-0E/β-18E.

.

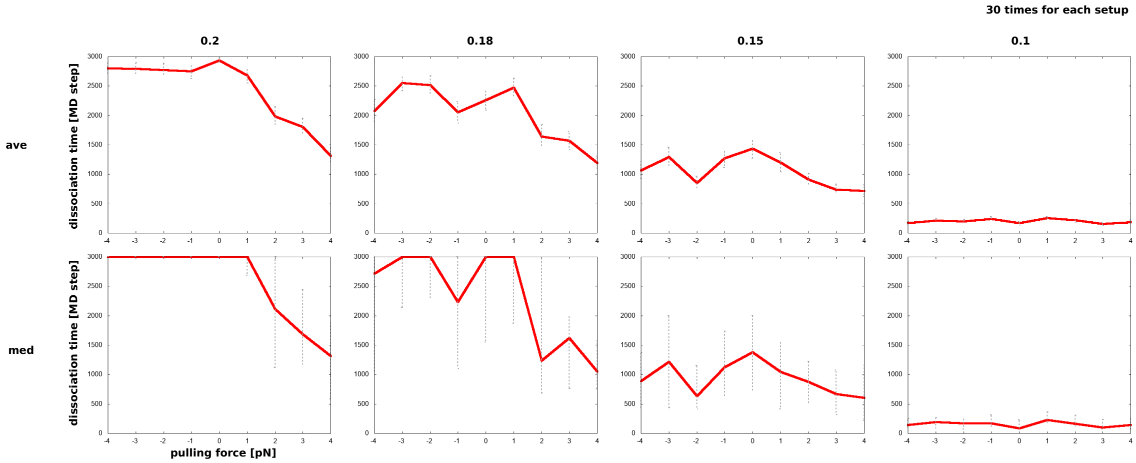


**Figure S6 High-affinity state dynein's dissociation time for each coefficient values.**
Average time it takes to dissociate for each external force in each coefficient. Thirty simulations were performed for each setting. The averages were taken as an additive average at the top and median at the bottom. Each error bar is the standard error and the interquartile range.

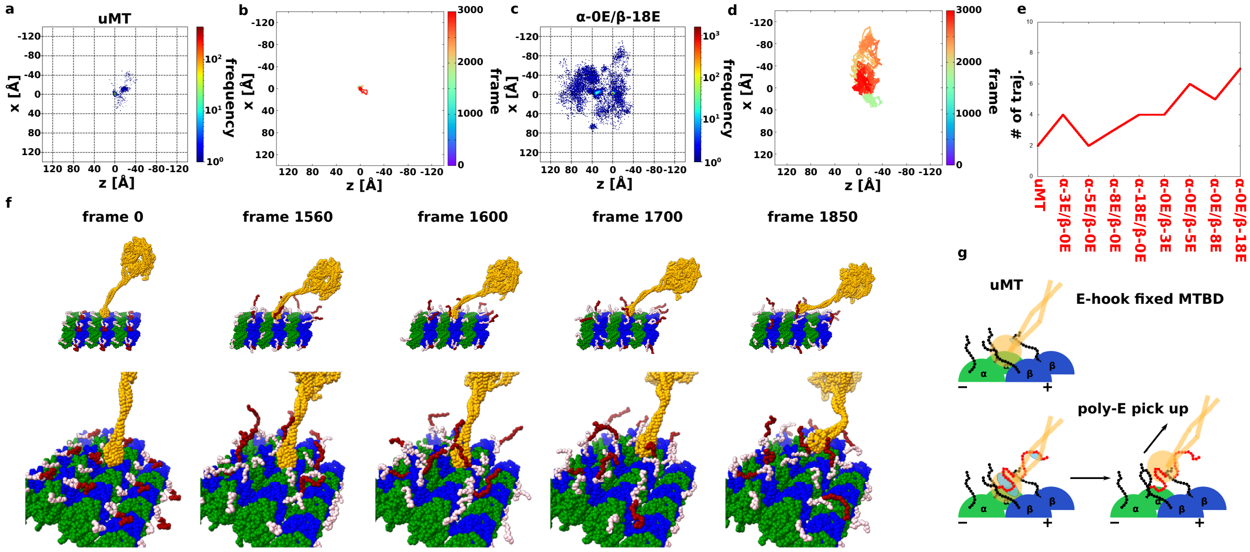


**Figure S7 High-affinity state dynein with poly-E MT.**

(a, c) Heat map of the position of the high-affinity state MTBD on the poly-E MT, with 10 trajectories overlaid. The coloring method is same with Fig. 1d. (a) is in the uMT case, and (c) is in the α-0E/β-18E case. (b, d) Representative trajectory of MTBD on the poly-E MT. The coloring method is same with Fig.1b. (b) is in the uMT case, and (d) is in the α-0E/β-18E case. (e) The number of trajectories in which the high-affinity state dynein in various poly-E environments moved from its initial position. Trajectories that moved more than 15 Å in each orientation from the initial position (0, 0) were counted as trajectories that moved. (f) Snapshots picked up from Fig.6c trajectory. The coloring method is same with Fig.1e. (g) Cartoon for understanding poly-E contact features. MTBD, α-tubulin, and β-tubulin colored orange, green, and blue. CTT is black, and poly-E is red. The contacting area of MTBD are circled in blue.

(a)

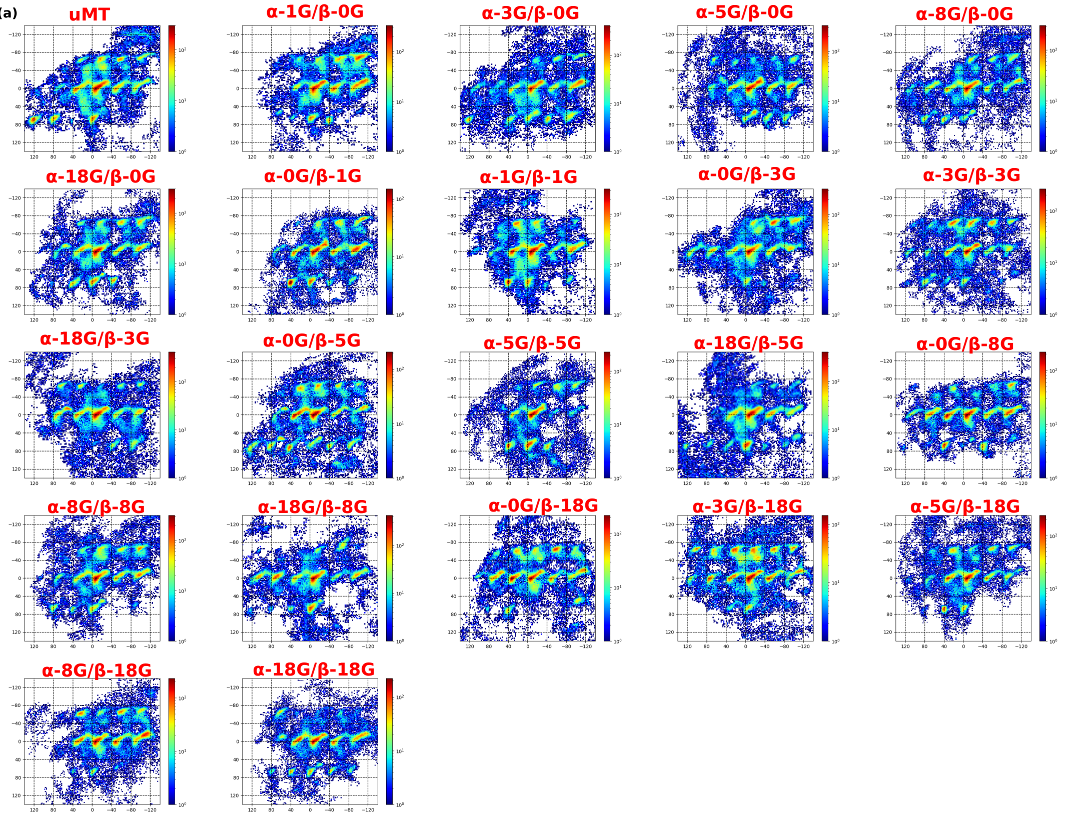
uMT
α-1G/β-0G
α-3G/β-0G
α-5G/β-0G
α-8G/β-0G
α-18G/β-0G
α-0G/β-1G
α-1G/β-1G
α-0G/β-3G
α-3G/β-3G
α-18G/β-3G
α-0G/β-5G
α-5G/β-5G
α-18G/β-5G
α-0G/β-8G
α-8G/β-8G
α-18G/β-8G
α-0G/β-18G
α-3G/β-18G
α-5G/β-18G
α-8G/β-18G
α-18G/β-18G

(b)

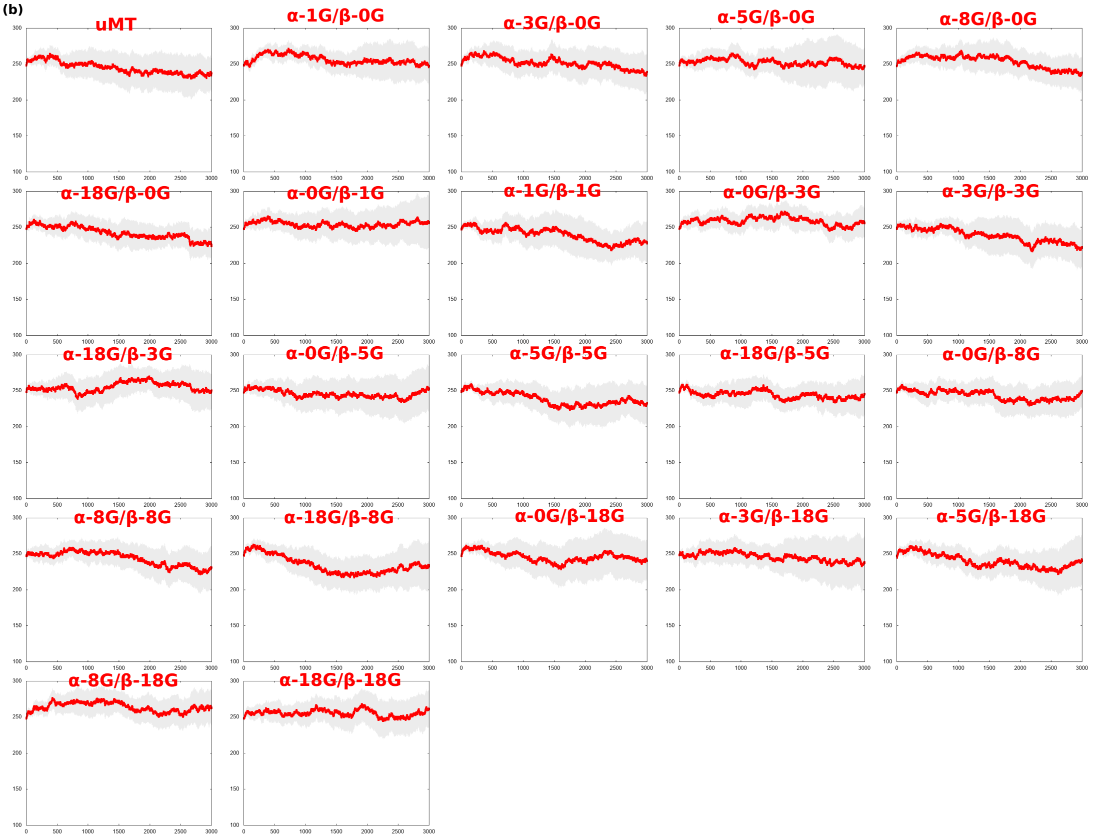
uMT
α-1G/β-0G
α-3G/β-0G
α-5G/β-0G
α-8G/β-0G
α-18G/β-0G
α-0G/β-1G
α-1G/β-1G
α-0G/β-3G
α-3G/β-3G
α-18G/β-3G
α-0G/β-5G
α-5G/β-5G
α-18G/β-5G
α-0G/β-8G
α-8G/β-8G
α-18G/β-8G
α-0G/β-18G
α-3G/β-18G
α-5G/β-18G
α-8G/β-18G
α-18G/β-18G

**Figure S8 MTBD position heatmap and direction between the linker and poly-G MT.**
(a) MTBD position heatmap for each setup. (b) Average distance between the tip of linker and the closest tubulin for each setup. Red line is the average of 20 trajectories for each frame, and gray is 95% confidence interval.

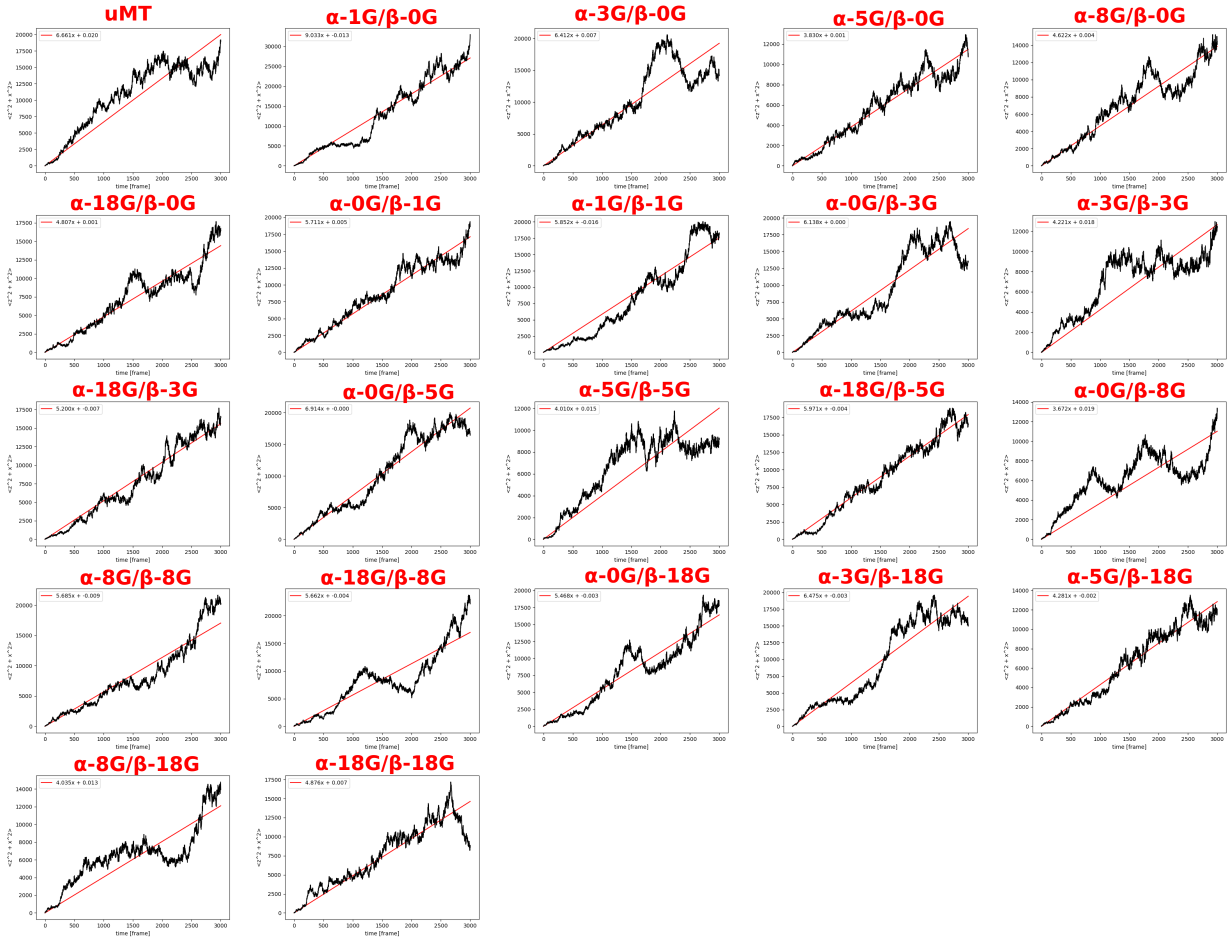


**Figure S9 MSD plot of the low-affinity state dynein on the poly-G MT.**
MSD plot for each frame (black) and its linear approximation line (red). The approximation is done by least-squares method.

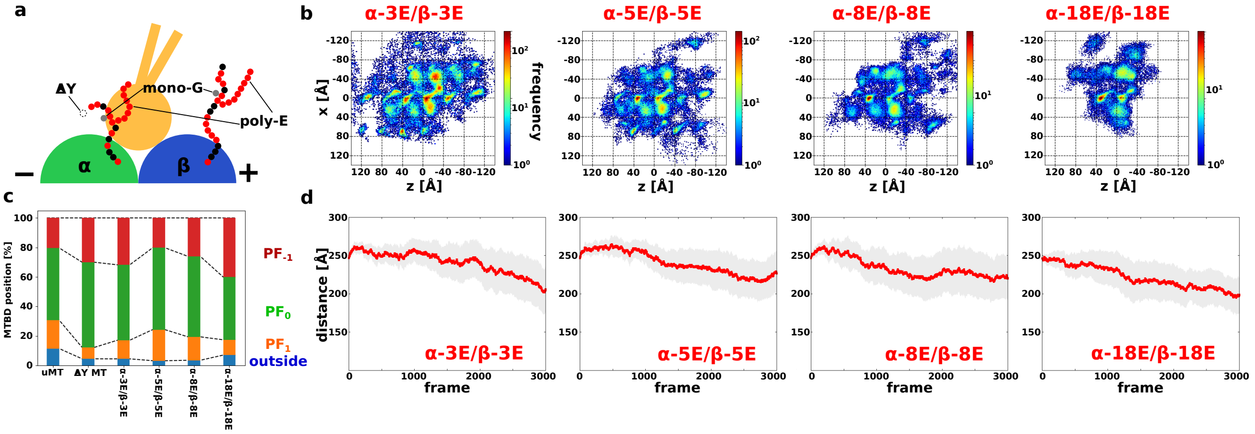


**Figure S10 Low-affinity state dynein with human tubulin code MT.**

(a) Cartoon for human tubulin code MT with detyrosination (ΔY), mono-G, and poly-E. (b) Heat map of the position of the low-affinity state MTBD on the human tubulin code MT, with 10 trajectories overlaid. The coloring method is same with Fig. 1d. The length of poly-E has several cases; three, five, eight, and 18. (c) Probability of localization of MTBD on the uMT, ΔY MT, several poly-E human tubulins code MT for each binding lane; $PF_{-1}$, $PF_0$, $PF_1$, and other percentages are red, green, orange, and blue, respectively. (d) The trajectory of the average direction between the tip of the linker to the closest tubulins in the several poly-E case.

### Supplementary Movies

**Movie S1 Representative trajectory for low-affinity state dynein with uMT.**
**Movie S2 Stepping mechanism cartoons for low-affinity state dynein with uMT.**
